# City life: airborne DNA metagenomic biodiversity monitoring reveals dynamic changes across time and space

**DOI:** 10.64898/2026.08.31.747859

**Authors:** Ned Peel, Piotr Cuber, Darren Heavens, Michael Giolai, Samuel Martin, Mark Alston, Darren Chooneea, Pia Aanstad, Raju Misra, Richard M Leggett, Matthew D Clark

## Abstract

Airborne environmental DNA can capture biodiversity across the tree of life, but low sample biomass makes rapid, untargeted detection technically challenging. We combined 45-min air collection, nanopore sequencing and real-time taxonomic analysis in a shotgun metagenomic workflow capable of producing results within 3 hours. Across 77 samples from 13 London sites, including a year of weekly sampling at the Natural History Museum Wildlife Garden, we detected 1,916 species spanning bacteria, fungi, plants and animals. Communities varied spatially and seasonally, shifting from plant dominance in spring to ascomycete dominance in summer and basidiomycete dominance in late autumn and winter. Plant read abundance increased with upwind vegetation, linking airborne signals to surrounding habitat. Detection of catalogued garden plants depended on reference availability, dispersal biology, plant size and proximity to the collector. Together, these findings establish airborne shotgun metagenomics as a platform for rapid, repeated and scalable biodiversity assessment across space and time.

## 1. Introduction

Biodiversity rates are now declining at a rate that is faster than previous mass extinction events **[Pereira et al. 2024]**. Preserving global biodiversity and ecosystems are United Nations (UN) Sustainable Development Goals **[UN 2020]**, which require biodiversity monitoring - ideally in real-time, to inform interventions and policy making. Environmental DNA (eDNA) is a suitably powerful and non-invasive technique for biodiversity monitoring **[Sahu et al. 2025]**. Metabarcoding of eDNA targeting one or more gene sequences is the most commonly used molecular tool to classify taxa presence, with low sample cost and high sensitivity, but struggles to be quantitative **[Shelton et al. 2022]**. Alternatively, shotgun metagenomic analysis of eDNA does not target specific gene loci, but instead sequences DNA fragments across the genomes of taxa present in an environmental sample, providing less biased and potentially more quantitative results than PCR-based methods **[Durand et al. 2025, Bell et al. 2021, Schmidt et al. 2022]**. The precision with which organisms can be classified is determined by methodology and their sequence similarity to sequences in public databases. For metabarcoding this is extensive, with over 1 million species in BOLD **[Ratnasingham et al. 2024]**. For shotgun analysis NCBI GenBank holds at least 581,000 formally described species **[Sayers et al. 2024]** with the Earth BioGenome Project (EBP) planning to generate high-quality reference genomes for 1.67 million eukaryotic species - its Phase 1 is sequencing at least one genome per eukaryotic family **[Blaxter et al. 2025]**.

The urban environment houses the majority of the world’s population, which is expected to rise to 68% by 2050 **[UN 2018]**. Urban areas are artificial constructs that present unique challenges to biodiversity monitoring e.g. large areas with limited flora and fauna interspersed with biodiversity rich parks, lakes and gardens containing native and imported species. London in the UK was the first city in the industrial age to reach a population of 1 million **[Census of Great Britain, 1801]**, with the choices of flora (e.g. avenue trees) shaped by the need to withstand the air pollution and deadly London smogs. With the switch from coal, new low emission zones and green technologies such as electric vehicles and renewable energy, pollution levels have dropped from their peaks **[Mayor of London 2022]**. Reduced pollution allows us to reimagine the urban environment to maximise function, health and well-being, with positive impacts for billions of people.

While metagenomic analysis is now a common technique, the majority of studies have been on soil or water samples, with air lagging in development because of the challenges it presents (for example, very low biomass). Air contains a wide variety of biological material including released plant pollen and fungal spores, aerosolised soil and water, dead cells, and naked DNA (see **[Berelson et al. 2025]** for recent review). Air is also the medium for many respiratory infections, which killed 4.25 million people a year even before the emergence of COVID-19 **[Mayor 2010]**. Recent studies have applied metabarcoding to airborne eDNA, including mitochondrial barcode primers targeting mammals **[Johnson et al. 2023]** and five metabarcodes targeting vertebrates (including mammals and birds), as well as arthropods, plants and fungi, using filters from an existing network of 15 UK heavy-metal monitoring sites **[Tournayre et al. 2025]**.

Previously we demonstrated that whole genome shotgun sequencing of air samples captures high metagenomic diversity and can be used as a surveillance tool for agricultural pathogens **[Giolai et al. 2024]**. While that study used Illumina short read sequencing, we also sequenced some of the same Illumina libraries on the then new (in 2015) Oxford Nanopore Technologies (ONT) MinION and found good concordance despite the lower accuracy and yields, but at the time the read count per sample was too low to ensure good sensitivity (Clark & Leggett pers. comm.). More recently **[Reska et al. 2024]** sequenced urban air using nanopore technology, noting their DNA extraction method underrepresents the harder to lyse gram-positive bacteria and fungal spore. Another study **[Nousias et al. 2025]** reported nanopore sequencing with a two day turnaround from the end of air sample collection to bioinformatics classification. However, their air collection period was up to 35 days, further increasing the reporting time.

Here we demonstrate that airborne DNA can provide a rapid and information-rich measure of biodiversity across a complex urban environment. We developed a shotgun metagenomic workflow combining short-duration air sampling with nanopore sequencing and taxonomic analysis, and applied it to samples collected across 13 sites in London and weekly over an entire year at the Natural History Museum (NHM) Wildlife Garden. The approach detected diverse bacteria, fungi, plants and animals, revealed marked seasonal turnover in airborne communities and identified taxa that were shared among, or specific to, individual sampling sites. Plant-derived reads were associated with surrounding vegetation, while comparisons with the well-characterised flora of the Wildlife Garden showed that taxon detection was strongly constrained by reference-sequence availability. Together, these results demonstrate the potential of rapid airborne shotgun metagenomics for repeated, broad-spectrum monitoring of biodiversity across space and time.

## 2. Results

### 2.1. Urban aerosol samples yield sufficient DNA for shotgun metagenomic sequencing

We wanted to assess whether our previous approach could be used to understand both the air microbial diversity within urban settings, and how similar the microbial communities present in different kinds of urban environments are. Since carrying out the agricultural work in **[Giolai et al. 2024]**, we moved from short read Illumina sequencing to long read nanopore technology, further refining our extraction and purification protocols (see Methods). We thus needed to confirm that DNA yields in urban settings were sufficient for this new approach. It has been previously shown that organisms differ in their ease of lysis, so we wanted to show that our approach was capable of extracting DNA across a broad range of organisms.

We took a Coriolis µ collector (Bertin, Montigny-le-Bretonneux, France) to 13 locations across London representing a diverse range of urban habitats (**Figure 1A**, **Supplementary Figure 1**) and collected air samples over 2 days (27th/28th August 2019). The same locations were visited 18 months later with an InnovaPrep Bobcat collector, with samples collected 5th-11th March 2021 (**Figure 1A, B**). Thus, we collected two sets of samples with two different air collectors at the same 13 sites, one set collected in summer, the other in spring. Hereafter, we refer to the August 2019 London Coriolis dataset as LC and the March 2021 London Bobcat dataset as LB. DNA was extracted, whole genome sequenced, and community composition determined. For the LC collections, DNA yields ranged from 3.10 ng (Vauxhall) to 19.36 ng (Marylebone) with a mean of 7.94 ng (**Supplementary Table S1**). Note, these figures exclude the Trafalgar sample (61.60 ng) which represented an outlier because it was contaminated with aphids. For the LB collections, DNA concentrations ranged from 1.66 ng (Natural History Museum wildlife garden) to 22.40 ng (Regents Park) with a mean of 6.34 ng.

**Figure 1:**
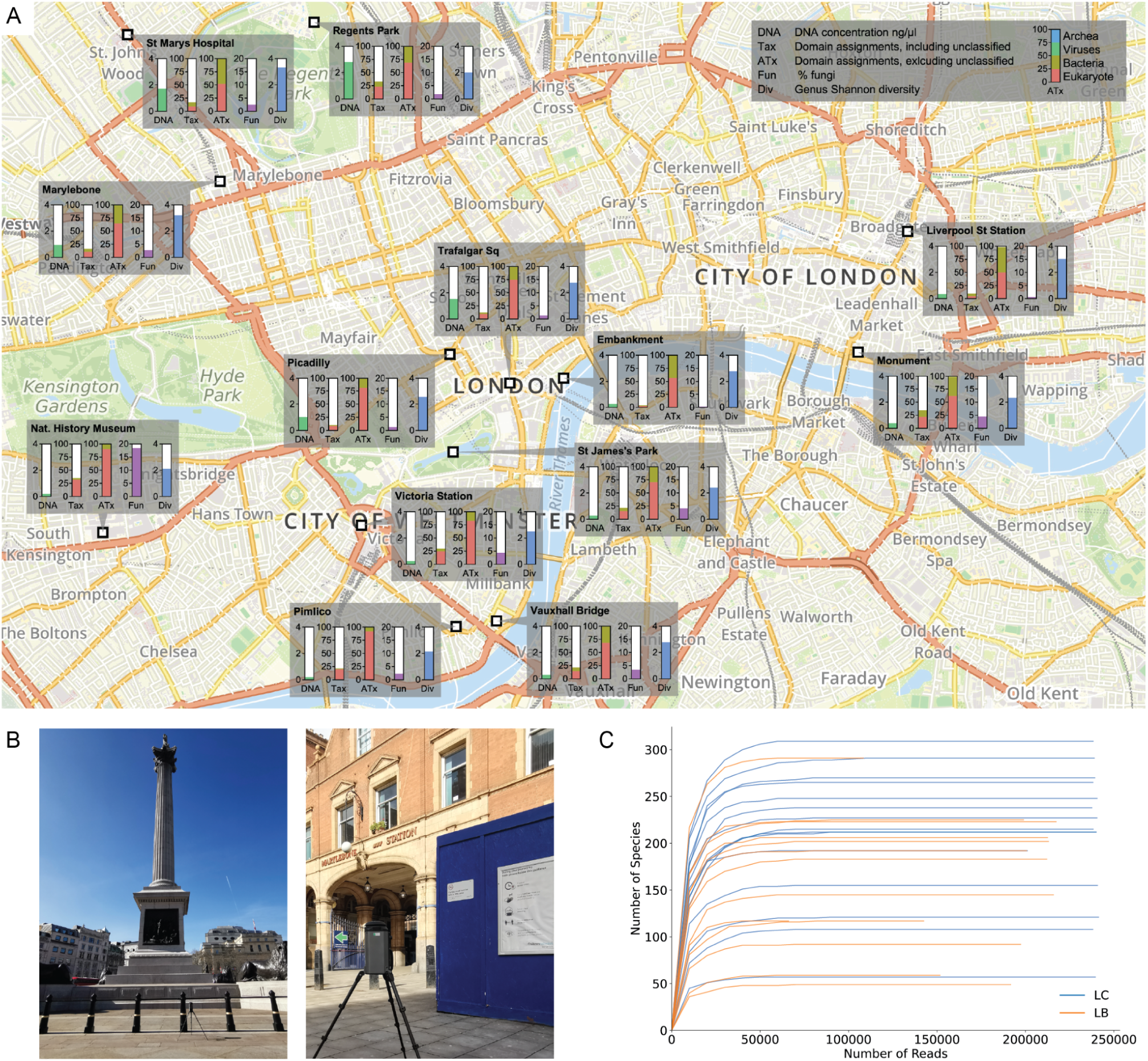
A) Map of London sampling locations. Each location is annotated with the DNA concentration, domain assignments (including unclassified and excluding unclassified), the percent of fungi, and the genus-level Shannon diversity from the LB dataset. B) Example collector placement (all locations in **Supplementary Figure 1**). C) Species accumulation curves for all samples from both the LC and LB datasets.

Sequencing the LC (August 2019) samples produced a mean of 1,922,105 reads per sample (range 902,111 - 4,095,964) with mean read N50 of 690 bp (range 603 - 841) (**Supplementary Table S2**). For the LB (March 2021) samples, sequencing produced a mean of 269,523 reads per sample (range 97,315 - 456,712) with mean read N50 of 1,913 bp (range 1,214 - 2,465) (**Supplementary Table S3**). Species accumulation curves (**Figure 1C**) plateaued, suggesting sequence data was sufficient to explore the sample diversity.

### 2.2. A diverse set of flora and fauna can be detected, with some common across all London sites and others unique to specific locations

All samples were analysed with a Lowest Common Ancestor (LCA) BLAST-based classification pipeline implemented in the MARTi tool [**Peel et al. 2025**]. Classification results can be viewed and interacted with at a project-specific MARTi instance which can be found at https://marti-air.cyverseuk.org/project/london-sites. Further analysis was carried out using custom scripts (see Methods). Across all LC (August 2019) samples and locations, the analysis identified 610 genera and 833 species, while for the LB (March 2021) samples, 432 genera and 639 species were identified. Combining both datasets resulted in identification of 812 genera and 1,177 species (**Figure 2A**). In the LC samples, species-level Shannon diversity ranged from 0.88 (Trafalgar Square) to 3.77 (Liverpool Street) with a mean value of 2.85 and a median of 2.99. For the LB samples, alpha diversity ranged from 2.06 (Regents Park) to 4.06 (St Marys Hospital), with a mean value of 2.74 and a median of 2.69 (**Table 1**).

**Figure 2:**
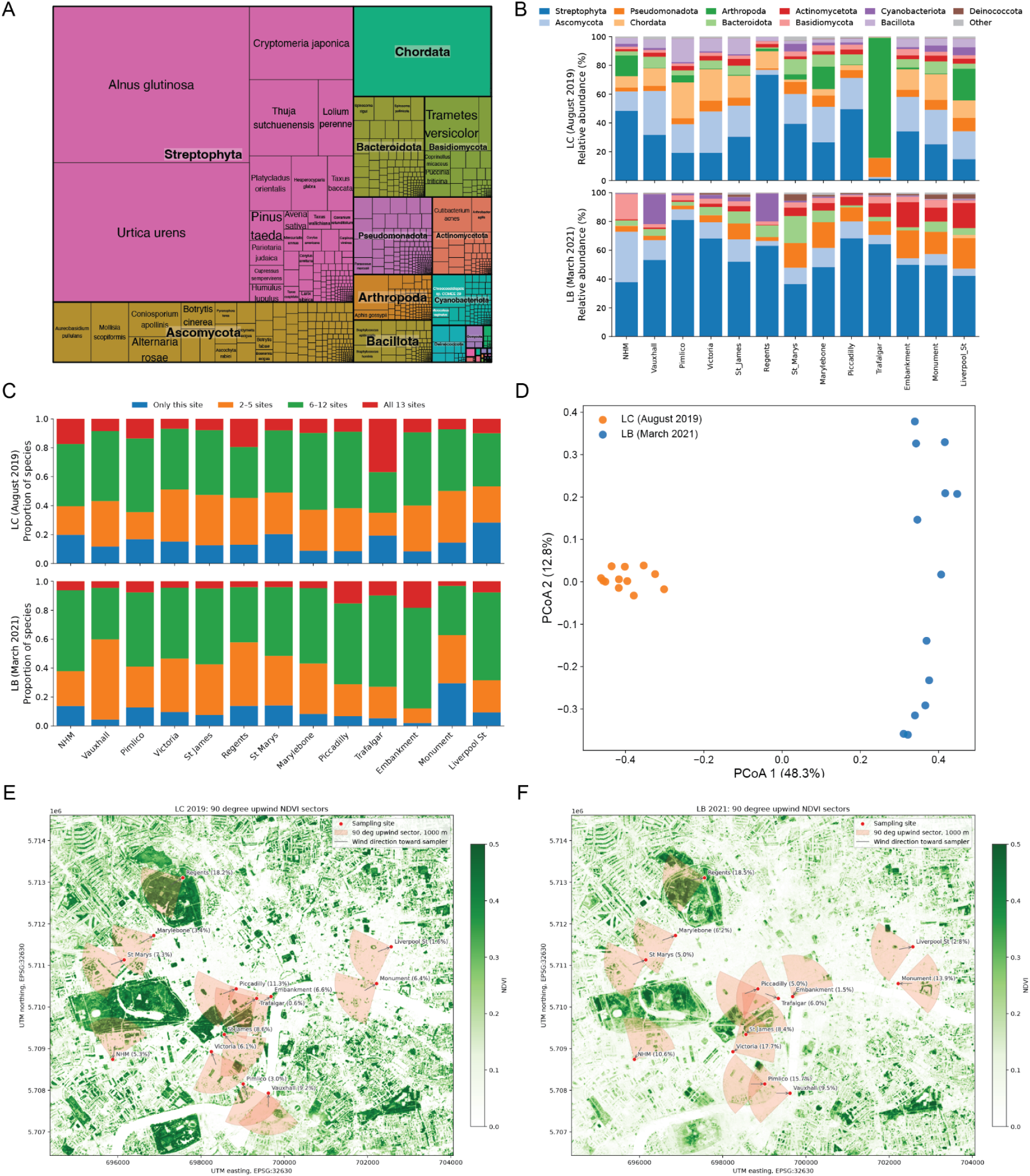
(A) Treemap of all 2019 and 2021 samples showing classified taxa grouped at phylum level. (B) Phylum level relative abundance plots for all locations and years. (C) Relative proportions of common and unique species across sites (only taxa with at least 0.005% abundance considered). Absolute counts can be found in **Supplementary Figure 3**. (D) Principal Coordinate Analysis (PCoA) of the air samples at species level (≥0.005% and ≥2 reads). The LC Trafalgar Square sample was excluded from panels A and D because the capture of an insect in the collector appeared to heavily bias the results (See **Supplementary Figure 2**). (E-F) Wind-aware NDVI source areas for London air-sampling sites. NDVI was calculated from Sentinel-2 Level-2A surface reflectance from 27 August 2019 for the LC sampling (E) and 9 March 2021 for the LB sampling (F). Red points mark air-sampling locations, with plant read relative abundance shown in brackets. Red shaded wedges indicate the upwind source area used for analysis: a 90-degree sector with a 1000 m radius. Figure contains Copernicus Sentinel data from 2019 and 2021. Wind observations obtained from Met Office MIDAS Open.

**Table 1:**
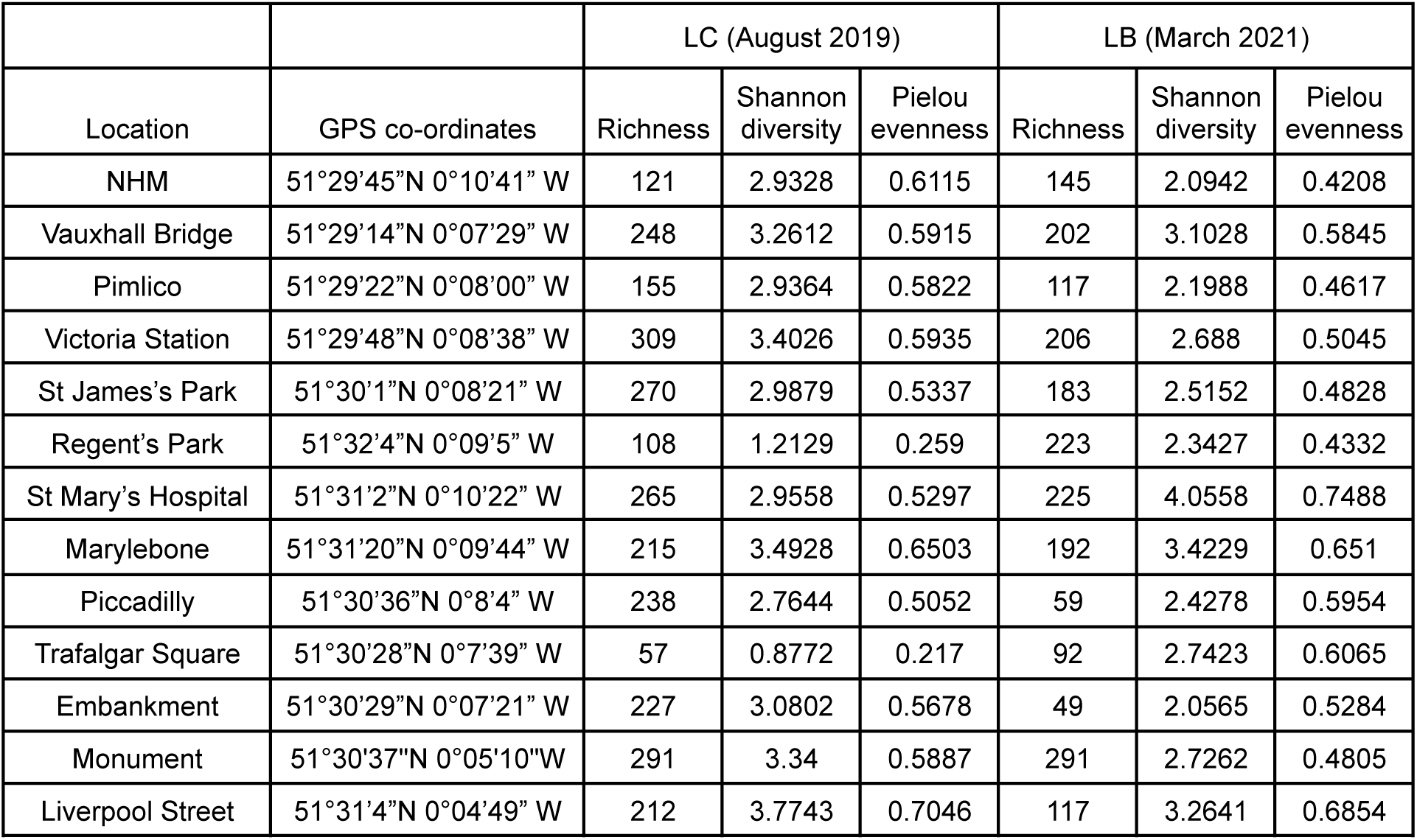
Species-level diversity at the collection sites. Species with abundance < 0.005% (of all analysed reads, classified and unclassified) are not considered.

A comparison of identified sequences across all locations demonstrated that there is a small core of species that was consistently detected across London. Considering taxa with at least 0.005% abundance, 21 species were detected at all 13 sites in the LC collection and nine in the LB collection. Each site also contained species not detected at any other sampled location, with the number of site-unique species ranging from 11 to 60 in LC and from 1 to 86 in LB (**Figure 2B,C**). In the LC samples, *Homo* (2.3% of all reads across all sites), and the fungal genera *Alternaria* (2.0%) and *Botrytis* (0.7%) were the most abundant genera across all sites. For the LB samples, the most abundant genera across all sites were *Alnus* (Alder trees, 5.6%), and the coniferous tree or shrub genera *Taxus* (1.2%) and *Thuja* (0.7%).

Principal Coordinate Analysis (PCoA) showed LC (August 2019) and LB (March 2021) samples clearly separated (**Figure 2D**), with one outlier (LC Trafalgar Square) removed due to the predominance of *Aphis gossypii* (we noted the presence of an insect in the collection tube, as the Coriolis has no insect screen). Separate per-collection PCoA plots are provided in **Supplementary Figure 2**. Normalised Difference Vegetation Index (NDVI) was positively associated with relative abundance of Streptophyta in both the LC and LB collections (**Figure 2E, F**). Using a 90-degree upwind sector with a 1000 m radius and a 1 h wind window, median upwind NDVI was correlated with Streptophyta read percentage in LB samples (Pearson’s r = 0.575, p = 0.040; Spearman’s r = 0.632, p = 0.021). The same relationship was also observed in LC samples after excluding Trafalgar Square (Pearson’s r = 0.695, p = 0.012; Spearman’s r = 0.462, p = 0.131), indicating that samples with greener upwind source areas tended to contain a higher proportion of plant reads.

### 2.3. Clear seasonal differences can be observed in the composition of the air microbiome

At one site, the NHM Wildlife Garden, sampling was carried out every week for an entire year (**Figure 3A**). Classification results can be viewed and interacted with at a project-specific MARTi instance which can be found at https://marti-air.cyverseuk.org/project/nhm-garden-weekly. Over the course of the year, the mean DNA yield per sample was 6.3 ng (range 0.9 to 72.2 ng) (**Supplementary Table S4**). Mean yield for spring, summer, autumn and winter were respectively 8.7, 2.2, 2.7, 10.7 ng. However, the winter yield was influenced by one particularly high yielding sample (72.2 ng) and without this, the mean would have been less than 4 ng. In contrast, the spring mean was less influenced by individual outliers and could reflect increased airborne biological material, for example from plant pollen. After sequencing and removal of short (<150 bp) and low quality (mean Q < 8) reads, there was a mean of 172,884 reads per sample (range 8,416 - 235,639) with mean read N50 of 2,555 bp (range 548 - 5,567) (**Supplementary Table S5**). Species accumulation curves approached plateaus, indicating that sequencing captured most of the diversity detectable under the applied thresholds (**Supplementary Figure 4A**). Across the year, 1,467 species were detected, with 927 detected in summer, 788 in spring, 763 in autumn and 620 in winter (**Figure 3C**; **Supplementary Figure 4B**).

**Figure 3:**
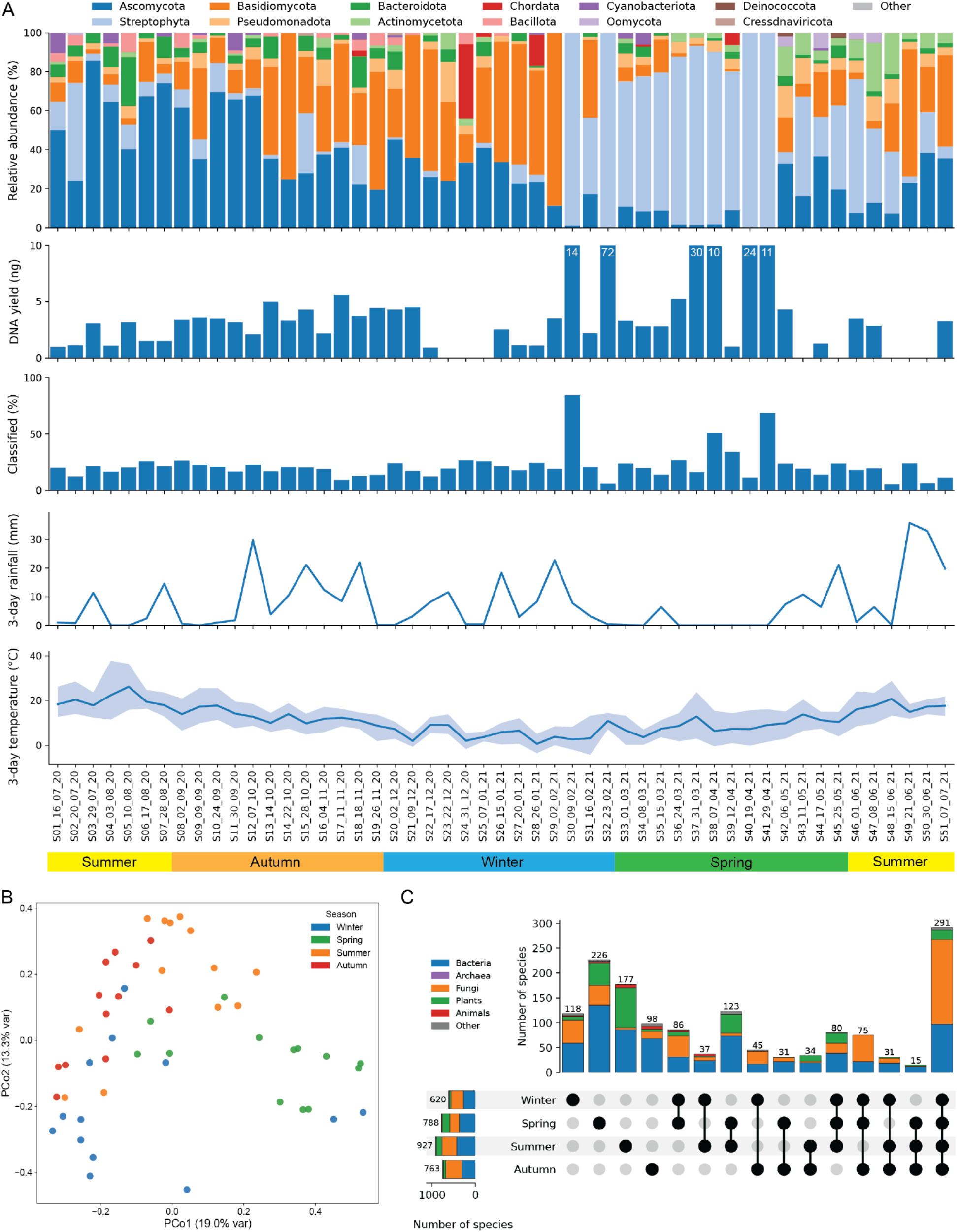
(A) Most abundant phyla read assignments for weekly samples (top); Percentage of reads that could be classified, DNA yield per sample and 3-day rainfall and average temperature. Seasons shown are UK meteorological seasons (based on calendar months). Where DNA yields exceed 10 ng, the bars have been capped and the yield written on the bar. (B) Principal Co-ordinate Analysis (PCoA) of the weekly samples, coloured by season. (C) UpSet plot showing overlap among species detected across seasons. Species were considered detected in a weekly sample if they had a relative abundance of at least 0.005% and were supported by at least two reads.

Clear seasonal differences can be seen in the taxonomic assignments (**Figure 3A**), with plants dominating in spring, Ascomycota dominating in summer and early autumn, and Basidiomycota dominating in late autumn and winter months. A principal co-ordinate analysis shows samples clustering roughly by season and with spring closer to summer and autumn closer to winter (**Figure 3B**).

Considering all weekly samples together, taxonomic assignments spanned bacteria, archaea, fungi, other microbial eukaryotes, plants, animals and viruses (**Figure 4**). Of the 1,467 species-level taxa meeting the detection criteria, 721 were bacterial, 451 fungal, 239 plant, 26 metazoan, 14 viral, 13 other eukaryotic, two archaeal and one an otherwise unclassified prokaryote. Ranked by mean relative abundance across all weekly samples, the bacterial profile was led by *Staphylococcus aureus* and included abundant soil- and dust-associated taxa such as *Pedobacter*, *Comamonadaceae*, *Chroococcidiopsis*, *Microlunatus*, *Friedmanniella*, *Pantoea*, *Pseudocnuella* and *Hymenobacter*. We also detect protists *Peronospora effusa* and *Albugo laibachii*, which are both plant pathogenic oomycetes (water moulds) common across the UK. Viral sequences represent a very small amount of classifications, but what we do recover are dominated by *Genomoviridae*, which have circular single-stranded DNA genomes that may be preferentially amplified during WGA. The two archaeal assignments, *Candidatus Nitrosocosmicus oleophilus* and *Methanosarcina mazei,* were also low abundance. Plant assignments were led by *Alnus*, *Carpinus* and *Quercus*, together with the conifers *Pinus* and *Cryptomeria*. These and the *Platanus* (plane trees) reflect the botanical landscaping typical of London. Metazoan assignments were less diverse and generally lower in abundance, consistent with animals not routinely releasing large quantities of airborne reproductive propagules analogous to pollen or fungal spores. Human sequences dominated the metazoan profile, while lower-abundance assignments included *Canis lupus* (dog), *Columba livia* (rock pigeon), *Mus musculus* (house mouse) as well as invertebrates such as *Myzus persicae* (Green peach aphid able to colonise many plant species).

**Figure 4:**
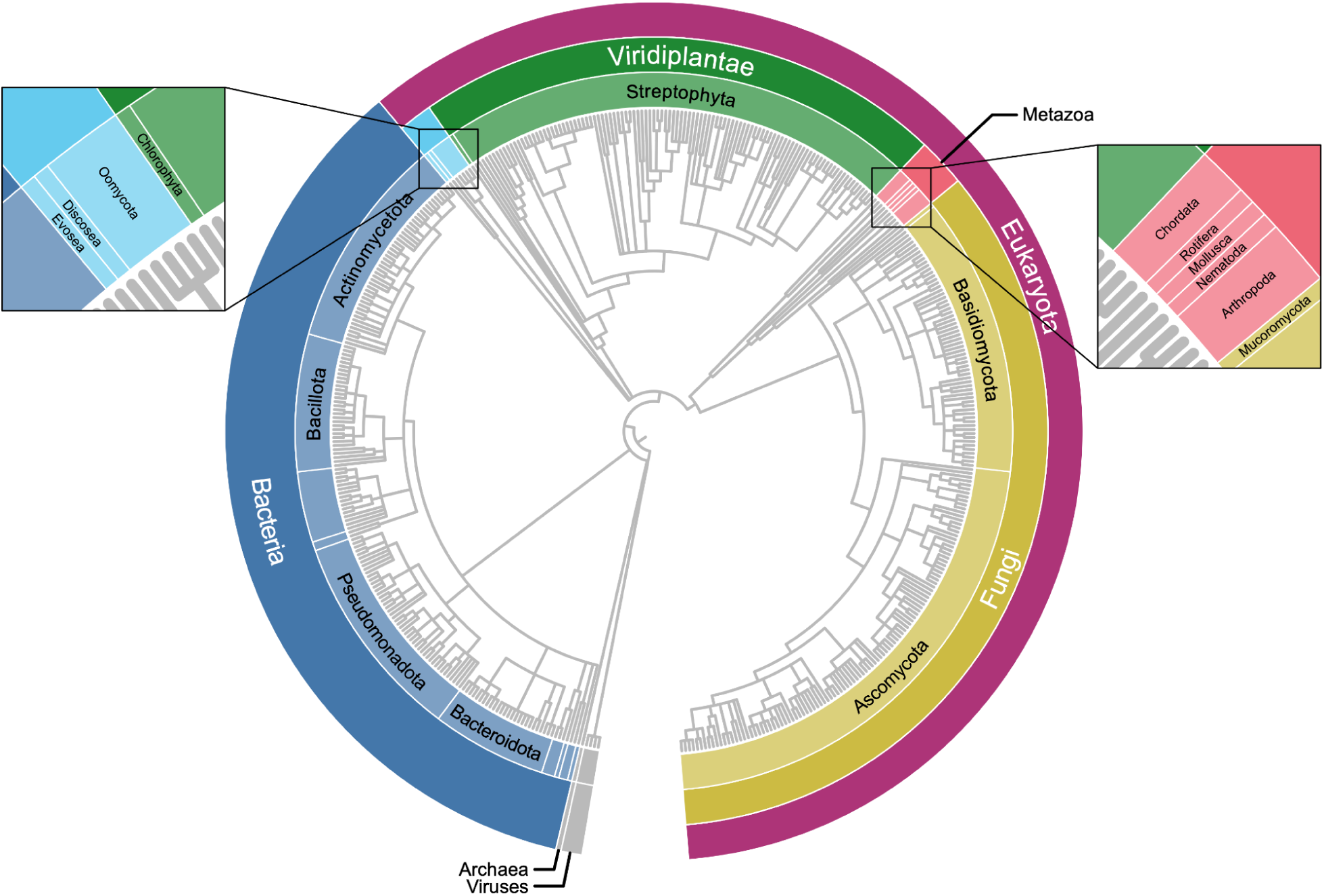
A high level view of the biodiversity found in the complete weekly sample set. The circular tree represents the taxonomic hierarchy of genera detected across all weekly samples using MARTi’s 0.1% minimum-support threshold. The size of each segment is proportional to the number of genera represented by it.

Fungi, principally Ascomycota and Basidiomycota, make up a high proportion of detected taxa from late summer through into winter. Classification with FUNGuild **[Nguyen et al. 2016]** showed a wide range of fungal guilds and growth forms detected throughout the year (**Supplementary Figure 6**). Guild composition was dominated by saprotrophic and plant-associated categories, while morphology was driven mainly by microfungus and polyporoid forms, with additional episodic contributions from agaricoid and yeast categories. These patterns were reflected in the detection of wood-decay and saprotrophic basidiomycetes such as *Stereum hirsutum*, *Trametes versicolor* and *Coprinellus micaceus*, together with plant-associated and microfungal taxa including *Aureobasidium pullulans*, *Mollisia scopiformis*, *Penicillium brevicompactum*, *Parastagonospora nodorum*, and *Botrytis cinerea*. Across seasons, the normalised plots indicate a shift from a stronger winter signal of wood-decay and undefined saprotrophic fungi towards a relatively greater contribution of plant-associated guilds in spring and summer, before returning to a more mixed assemblage in autumn. Morphologically, polyporoid fungi made up a large proportion of fungal abundance in winter, whereas microfungal forms contributed a larger proportion in spring and summer. These patterns are consistent with the broader seasonal taxonomic trends in the dataset, in which Ascomycota dominate summer and early autumn samples and Basidiomycota dominate late autumn and winter.

MARTi can identify antimicrobial resistance (AMR) genes in sequence data using the CARD database (Comprehensive Antibiotic Resistance Database **[Alcock et al. 2023]**). An analysis of the weekly samples from the Wildlife Garden identified 2,557 hits to 42 different AMR genes at ≥ 90% identity match over the course of the year (**Supplementary Table S6**). The vast majority of these (98%) were found in samples collected between September and February (**Supplementary Table S7**). MARTi attempts to identify host bacteria for AMR genes by using flanking sequence and in these data it placed the majority of AMR hits (70%) within Staphylococcus. An analysis of the correlation between taxa and AMR hits unsurprisingly showed strong correlation with *Staphylococcus aureus* (Spearman’s ρ = 0.771, FDR-adjusted q = 3.92 × 10⁻⁸), as well as several other bacteria including *Listeria monocytogenes* and *Limosilactobacillus fermentum* (**Supplementary Figure 5**).

### 2.4. Species detection is strongly influenced by reference availability and proximity to source

We looked at the samples from the Natural History Museum Wildlife Garden, an area of approximately one acre used for recreating several UK habitats (**Figure 5A, B**). Since sampling took place, this area has been redeveloped into the Nature Discovery Garden with a much changed set of taxa. As the flora in this area were well catalogued (see **Supplementary Table S8**) their presence/absence in the air sequencing data can be compared and used to see what affects our method’s ability to detect known species in a sampling area.

**Figure 5:**
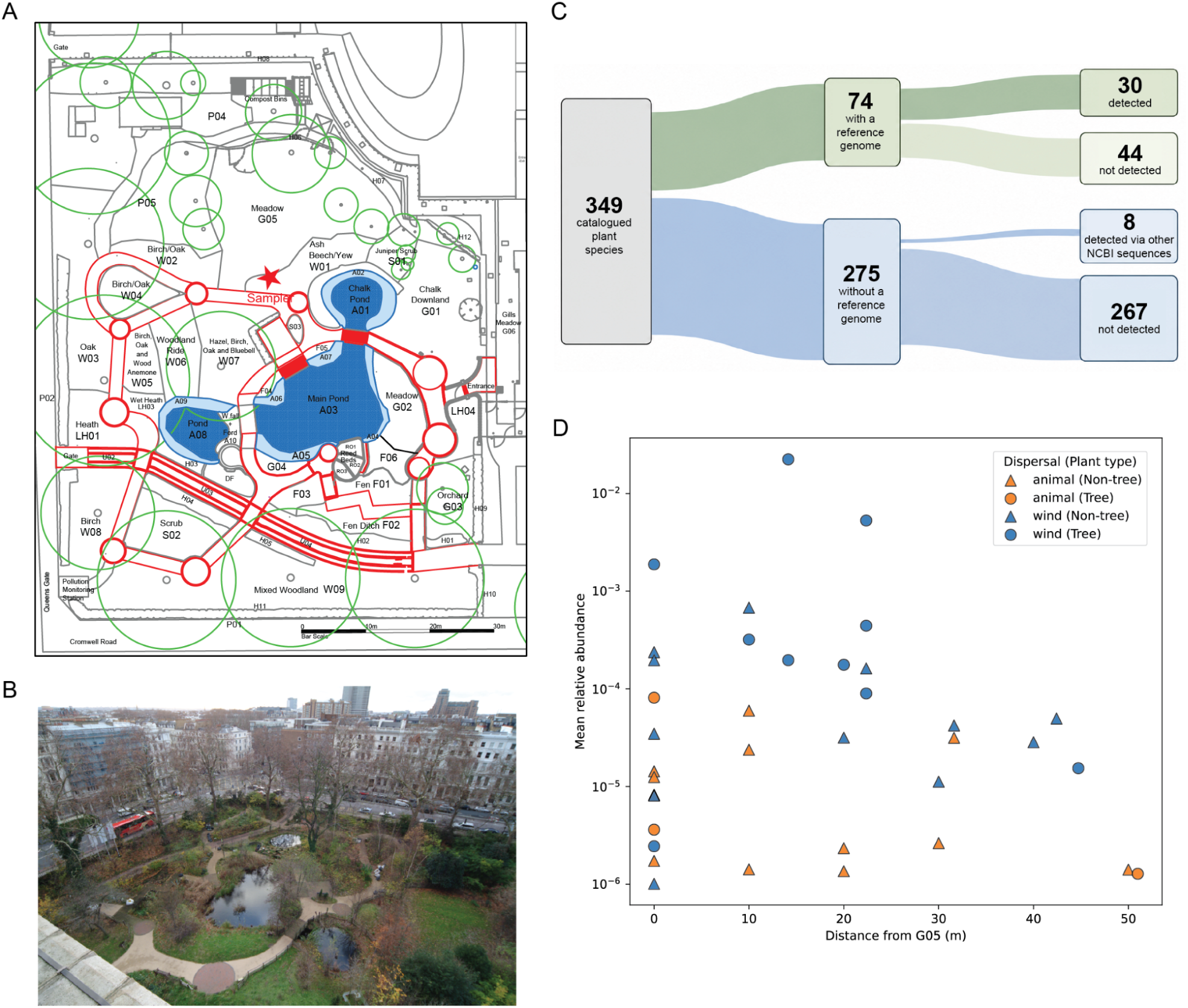
(A) Map of NHM wildlife garden, with collector placement shown with a star in Meadow G05 towards the top centre. (B) Aerial view of the wildlife garden, taken in 2009 but representative of site at time of sampling. (C) Sankey plot illustrating the catalogued plant species, the number with reference genomes and the number detected. (D) Mean relative abundance of plant taxa detected by our pipeline and known to be present in the garden, plotted as a function of distance (m) from the Meadow, G05. Map and photograph © The Trustees of the Natural History Museum, London. All Rights Reserved.

Of the 349 plant species known to be present in the garden, 74 had genome assemblies available at NCBI when the nt database used in our analysis was released (**Supplementary Table S9**). We detected 30 of these 74 (40.5%), compared with eight of the 275 species without an available genome assembly (2.9%); these eight had other, non-genome sequences represented in NCBI. In total, 38 garden species were detected across the year of sampling (**Figure 5C**). At the broader genus level, we detected 63 of the 228 genera represented in the garden inventory (27.6%), including 25 genera not recovered through species-level matching.

We hypothesised that wind-pollinated species would be more readily detected in collected air than non-wind-pollinated species. Consistent with this, among the 74 Wildlife Garden plant species with genome assemblies available in NCBI, 14 of 24 wind-pollinated species were detected compared with 16 of 50 animal-pollinated or mixed/other species, corresponding to detection rates of 58.3% and 32.0%, respectively. When all 38 detected garden plant species were considered, including the eight species detected through non-genome NCBI sequences, wind-pollinated or wind/spore-dispersed taxa accounted for 63.5% of detected plant species and 98.6% of plant reads. There was also evidence that signal decreased with distance from the collector, particularly among detected wind-pollinated species (**Figure 5D**). The prevailing wind direction is also likely to have played a part in determining the potential for detection of plant species, as pollen and cells from some plants would be blown away from the collector.

## 3. Materials and Methods

### 3.1. Sample capture

For the geographical dataset across London (‘L’),13 samples were collected from the same locations in 2019 and 2021 (**Figure 1**, **Figure 2**). The 2019 samples were collected using a Coriolis µ collector (Bertin, Montigny-le-Bretonneux, France) with continuous air intake mode at 300 L/min, 30 minute collection time (9,000 L total). The 2021 samples were collected using an InnovaPrep Bobcat collector (Innovprep, Drexel, Missouri, USA) in continuous air intake mode at 200 L/min, for 45 minute collection time (9,000 L total). Sample location codes were consistent between 2019 and 2021, except ‘C’ (Coriolis in 2019) and ‘B’ (Bobcat in 2021). Each sample was obtained from processing a total air volume of 9000 litres (**Table 2**). Average temperature and wind speed values were calculated as the mean of readings from the beginning, middle and end of air-sampling (**Table 2**) at the same level as the air collector (approximately 1.5 m). The measurements were recorded using a Testo 410-2 thermo-anemometer (Testo, Alton, Hampshire, UK). GPS coordinates were taken from the Apple Compass app (iPhone) or the GPS Coordinates app (Android).

**Table 2:** Sample site environmental metadata; temperature, humidity, wind speed and volume of air collected. *Battery ran out on LC7 (St James), hence the lower volume of air collected.

|  | August 2019 (LC) |  |  |  | March 2021 (LB) |  |  |  |
| --- | --- | --- | --- | --- | --- | --- | --- | --- |
| Location | Temp. (C) | Humidity (%) | Wind speed (m/s) | Total air volume collected (L) | Temp. (C) | Humidity (%) | Wind speed (m/s) | Total air volume collected (L) |
| NHM | 26 | 54 | 1 | 9000 | 11.9 | 38.4 | 0.25 | 9000 |
| Vauxhall Bridge | 32 | 36 | 0.26 | 9000 | 12.4 | 49.2 | 0.17 | 9000 |
| Pimlico | 33 | 34 | 0.26 | 9000 | 13.2 | 41.9 | 0.33 | 9000 |
| Victoria Station | 32 | 36 | 0.26 | 9000 | 11.2 | 46.1 | 1.53 | 9000 |
| St James's Park | 32 | 36 | 0.26 | 4500* | 11.9 | 47.0 | 0.57 | 9000 |
| Regent's Park | 24.38 | 60.17 | 1 | 9000 | 13.7 | 50.3 | 0.77 | 9000 |
| St Marys Hospital | 25.93 | 54.85 | 1 | 9000 | 13.3 | 54.3 | 1.40 | 9000 |
| Marylebone | 25.16 | 47.3 | 1.3 | 9000 | 14.6 | 47.9 | 0.77 | 9000 |
| Piccadilly | 26.47 | 39.19 | 1.5 | 9000 | 8.4 | 59.7 | 0.57 | 9000 |
| Trafalgar Square | 28.45 | 36.44 | 1.3 | 9000 | 15.9 | 36.7 | 0.50 | 9000 |
| Embankment | 32.26 | 28.32 | 2.1 | 9000 | 10.2 | 45.2 | 0.4 | 9000 |
| Monument | 26.45 | 38.81 | 0.8 | 9000 | 10.0 | 48.6 | 0.93 | 9000 |
| Liverpool Street | 24.46 | 46.76 | 4.3 | 9000 | 12.5 | 46.5 | 0.13 | 9000 |

Samples for the weekly dataset were collected in NHM’s Wildlife Garden, South Kensington, London, UK (GPS coordinates: 51.496085, −0.177891 and **Figure 1A**). Weekly sampling was across all four seasons, starting on 16th July 2020 and ending 7th July 2021. An InnovaPrep Bobcat collector was used in continuous air intake mode at 200 L/min, for 45 minute collection time.

### 3.2. DNA extraction

#### Sample concentration

Liquid sample, either direct from the collection vessel (Coriolis), or eluted from the air filter using the manufacturers buffer (InnovaPrep), were then filtered using 0.2 µm PVDF membranes in Swinny holders (both MilliporeSigma, Gillingham Dorset, UK). Samples on these filters were stored at −20 °C until further processing.

#### Lysis and extraction

The Swinny filter was placed in a DNeasy PowerSoil Pro Kit 2 ml tube and 100 µl of CD1 solution and 250 mg of (beating) beads (Qiagen, Manchester, United Kingdom) were added. This was bead beaten in a TissueLyser II (Qiagen) for 5 mins at a speed setting of 25 Hz. Tubes were centrifuged for 1 minute at 13,000 rpm, and the supernatant transferred to a 1.5 mL tube. A 1x AMPure XP (Beckman Coulter, Amersham U.K.) bead clean-up was performed (100 µl AMPure XP beads, followed by 2x wash in 200 µl 70% EtOH) and eluted in 8 µL of DNase/RNase free water for 5 minutes. DNA extractions were quantified using Qubit dsDNA HS reagents.

#### Controls

Two negative controls SN1 and SN2 (molecular grade water, MilliporeSigma) were also processed throughout the whole pipeline alongside true samples.

#### DNA pipeline sample processing

#### Whole Genome Amplification (WGA)

We used a reduced volume reaction for the Repli-G WGA kit (Qiagen), combining 3.75 µL DNA with 0.25 µL Repli-G DLB buffer, incubated for 3 minutes to denature the gDNA extracted. Then 0.4 µL of Repli-G Ultrafast Stop solution was added to the reaction and mixed well. This was followed by 16 µL of Repli-G Ultrafast reaction buffer and 1 µL of Repli-G Ultrafast polymerase, mixed well and incubated for 90 minutes at 30 °C. A 1x AMPure XP bead clean-up followed (20 µl AMPure XP beads, 2x wash in 200 µl 70% EtOH) and was eluted in 17 µL of DNase/RNase free water (MilliporeSigma).

#### Endonuclease debranching treatment

WGA is highly branched and can clog nanopores, so we debranched to maximise sequence yields. We added 4 µL 5x S1 nuclease buffer and 1 µL (100 U/µL) S1 nuclease (Thermo Scientific) to the clean WGA reaction and incubated for 15 minutes at 37 °C. After this, we performed a final clean-up step with 1x AMPure XP bead clean-up (20 µl AMPure XP beads, 2x wash in 200 µl 70% EtOH) and eluted in 9 µL 1x TE buffer. Reactions were quantified using Qubit Broad Range reagents. In cases where DNA concentration post endonuclease treatment was low, or if it was necessary to repeat sequencing and we did not have enough post-endonuclease treatment samples left, WGA and S1 treatment were repeated using leftover DNA extraction.

### 3.3. Library construction and sequencing

DNA library preparation and sequencing followed Oxford Nanopore Technologies’ (ONT’s) protocol for Rapid Barcoding using the Rapid Barcoding Kit (SQK-RBK004) MinION R9.4.1. (FLO-MIN106D) hereafter called “R9”. Libraries were constructed from 50 ng of WGA and debranched material. Up to 12 libraries were pooled together for MinION sequencing, with MinKNOW using the latest basecaller and demultiplexing (this has been subsequently repeated with newer software, see below), we targeted >50,000 reads per sample.

### 3.4. Bioinformatics

#### Basecalling and demultiplexing

Raw FAST5 files were converted to the newer POD5 format using pod5 v0.3.6. Basecalling and demultiplexing were performed with Dorado v0.6.2 using the super-accuracy model dna_r9.4.1_e8_sup@v3.6.

#### Read statistics and subsampling

Per-sample read statistics, including total read count, mean read length, N50, and mean quality score, are provided in **Supplementary Tables S2, S3 and S5** for the LC, LB and weekly datasets, respectively. To standardise sequencing depth across samples, reads were randomly subsampled to a maximum of 250,000 reads per sample. Samples yielding fewer than 250,000 reads were retained in full.

#### Taxonomic classification

Basecalled reads were analysed using MARTi v0.9.19. MARTi’s internal prefilter retained reads ≥150 bp with a minimum mean quality score of Q8 prior to taxonomic classification. Taxonomic assignment was performed using MARTi’s BLAST-based **[Camacho et al. 2009]** Lowest Common Ancestor (LCA) pipeline against the NCBI nt database (March 2024 release). BLAST searches were conducted with DUST low-complexity filtering enabled (-dust 15 64 1; **[Morgulis et al. 2006]**). Alignments were retained for LCA consideration if they met the following criteria: (i) minimum sequence identity of 75%, (ii) minimum alignment length of 150 bp, and (iii) bit score ≥90% of the highest-scoring alignment for that read. The LCA for each read was computed from all retained alignments passing these thresholds.

#### Relative abundance and taxon filtering

Relative abundance was calculated as the number of reads assigned to a taxon divided by the total number of reads analysed in that sample, including classified and unclassified reads. A minimum relative abundance of 0.005% was used for most species-level analyses. Where an additional requirement for at least two supporting reads was applied, this is stated for the relevant analysis below.

#### Accumulation curves

Species and genus accumulation curves were derived from MARTi’s real-time chunked taxonomic tree JSON files. For each sample, taxa were first filtered using the final chunk only, retaining taxa supported by ≥2 reads and representing ≥0.005% of the total analysed reads. The set of taxa were then tracked retrospectively across earlier chunks to determine the cumulative number detected at each sequencing depth.

#### Principal coordinates analysis

Beta-diversity between samples was assessed using Bray-Curtis dissimilarity and visualised by principal coordinates analysis (PCoA). For each sample, taxonomic tables were derived from the MARTi tree JSON files and filtered independently to retain taxa meeting the minimum abundance threshold (≥2 reads and ≥0.005% of total analysed reads). Pairwise Bray-Curtis dissimilarities were computed between samples, and PCoA was performed by eigen-decomposition of the centred distance matrix. For the LC versus LB comparison, samples were coloured by collection. For the weekly analysis, samples were coloured according to meteorological season (Winter: December-February; Spring: March-May; Summer: June-August; Autumn: September-November).

#### Seasonal overlap analysis

Species-level assignments were obtained from the final MARTi taxonomic table for each weekly sample. A species was considered detected in a sample if it represented at least 0.005% of all analysed reads and was supported by at least two reads. Samples were assigned to meteorological seasons according to their collection date. A species was considered present in a season if it was detected in at least one sample assigned to that season. Seasonal sets and their intersections were visualised using UpSetPlot v0.9.0 **[Lex et al. 2014]**, with intersection sizes subdivided by broad taxonomic group.

#### Treemap visualisation

Species-level assignments from MARTi were filtered per sample to retain taxa supported by ≥0.005% of total analysed reads. For each species, mean relative abundance was computed across the selected sample set. Species were grouped by NCBI lineage at phylum level and visualised as a treemap using the R package treemap v2.4-4 **[Tennekes, 2023]**, with rectangle area proportional to abundance and colours assigned by phylum.

#### Circular tree

Genus-level lineages were extracted from MARTi taxonomic trees generated using a minimum-support threshold of 0.1% of taxonomically assigned reads within each sample. NCBI identifiers were pooled across samples and deduplicated, and their complete NCBI taxonomic lineages were converted into a rooted Newick hierarchy. The hierarchy was displayed in circular format using Interactive Tree of Life (iTOL) v6 **[Letunic and Bork 2024]**.

#### Fungal functional annotation

Fungal taxa reaching at least 0.005% relative abundance in one or more weekly samples were annotated using FUNGuild v1.1 **[Nguyen et al. 2016]**. Matched annotations from all confidence rankings were retained. Weekly guild and growth-morphology abundances were calculated by summing the relative abundances of assigned taxa. Seasonal mean abundances were normalised across matched fungal taxa within each season. Categories contributing at least 1% of pooled weekly abundance or 5% in any week were shown separately, with the remainder grouped as “Other”. Taxa with multiple annotations contributed their full abundance to each category, so seasonal totals could exceed one.

#### AMR identification and host assignment

Antimicrobial resistance genes were identified from the weekly samples using MARTi with the Comprehensive Antibiotic Resistance Database (CARD) **[Alcock et al. 2023]**. AMR read abundance was defined as the number of supporting reads, and AMR gene richness as the number of distinct retained genes in each sample. MARTi’s walkout analysis was used to infer putative bacterial hosts by taxonomically classifying sequence flanking the CARD-aligned region. Host assignments were therefore treated as putative rather than definitive associations.

#### Meteorological data

Daily rainfall and air temperature observations were obtained from the UK Met Office MIDAS Open dataset for a nearby meteorological station (Heathrow) and used to characterise local weather conditions during the NHM Wildlife Garden weekly sampling period (2020–2021). For each sampling date, meteorological variables were summarised over the three calendar days preceding the collection date. Three-day rainfall was calculated as cumulative precipitation (mm) across this period, and three-day temperature metrics comprised the minimum, maximum and mean air temperature over the same interval.

#### Satellite data

NDVI was calculated from Sentinel-2 Level-2A surface reflectance using the 10 m red (B04) and near-infrared (B08) bands as (B08 - B04) / (B08 + B04). Imagery from 27 August 2019 was used for the LC collection, and 9 March 2021 was used for the LB collection. To account for wind direction during sampling, hourly mean wind observations from Met Office MIDAS Open at Battersea Heliport were matched to each sample using the hourly record falling within the 1 h window ending at sample completion. For each sample, NDVI was summarised within a 90-degree upwind sector with a 1000 m radius, centred on the mean wind direction towards the sampler. Median NDVI within this sector was used as the wind-aware vegetation metric. Associations between median upwind NDVI and Streptophyta read percentage were assessed separately for LB and LC using Pearson and Spearman correlations; the Trafalgar Square sample was excluded from the August 2019 (LC) correlation analysis.

#### NHM Wildlife Garden

Taxa detected in weekly air samples from the NHM Wildlife Garden were compared with vascular plant records for the garden for 2020 and 2021. Species were considered detected if they met the 0.005% minimum abundance threshold defined above in at least one sample. Mean relative abundance was calculated across all weekly samples, including zeros for samples in which the species was not detected. Where multiple records existed for a species within the NHM garden, the minimum recorded distance to G05 was used. Genome assembly availability was assessed using NCBI RefSeq and GenBank assembly summary files. Assemblies were considered available if released before the nt database build date (March 5, 2024) and not withdrawn prior to this date. Species were classified as having genome representation if at least one qualifying assembly was associated with their NCBI taxonomy identifier.

## 4. Discussion

Here we describe a new improved eDNA approach for airborne biological material which combines optimised metagenomic shotgun nanopore sequencing and bioinformatics to monitor biodiversity in a variety of locations and habitats (including urban) and can generate analysed results within 3 hours end-to-end (45 mins air collection, <60 min molecular biology, <60 min sequencing and analysis). Rapid analysis is achieved using Metagenomics Analysis in Real Time (MARTi) **[Peel et al. 2025]** which takes advantage of the nanopore sequencing platform’s capability for progressive analysis - though speed may be limited by the choice of classification algorithm, database and compute availability (e.g. BLAST vs nt may not be possible in under an hour). Our pipeline can filter biological material from 9,000 litres of air to generate 8 µl of final DNA, over a billion fold concentration. Other studies have conducted nanopore shotgun sequencing of air, but either they used a longer (7-35 day) sampling time to obtain enough DNA, or sometimes reported a bias in detected taxa **[Reska et al. 2024, Nousias et al. 2025]**. Faster results are useful in most fields, especially public health, but also for biodiversity monitoring where eDNA analysis can inform researchers of unseen but present organisms.

Our analysis of 77 samples from 13 sites identified 1,916 species and 1,149 genera across all kingdoms of life (using a per-sample detection threshold of at least 0.005% abundance and two reads) and revealed both seasonal and geographical variation in airborne eDNA across London. This level of taxonomic richness is comparable to that reported by **[Tournayre et al. 2025]**, who recovered 1,556 amplicon sequence variants assignable to unique taxa, representing 1,220 genera, from 185 samples across 15 UK sites using metabarcoding. We detected microbial species, including harder-to-lyse fungi and Gram-positive bacteria, as well as plants and animals, with wind-pollinated plants often dominating samples during their flowering periods.

Plant pollen is the principal allergen for hundreds of millions of people with hay fever, whose symptoms vary with season, location, weather and pollution and can progress to asthma **[Savouré et al. 2022]**. Our results suggest that the speed of this approach could support daily, species-level pollen monitoring and potentially identify allergen-associated allelic variants, enabling airborne exposure profiles to be compared with symptoms. Across the complete sampling year, wind-pollinated or spore-dispersed taxa accounted for 98.62% of plant-assigned reads but only 63.5% of detected plant species, whereas animal-pollinated taxa accounted for 1.38% of reads but 36.5% of species. Thus, although wind-dispersed material dominated read abundance, the method still detected substantial species diversity among plants that do not primarily disperse pollen by wind. Read proportions are, however, compositional and should not be interpreted directly as estimates of organismal or particle abundance. Although shotgun sequencing can be more quantitative than PCR-based metabarcoding, read abundance remains influenced by source biomass, particle and cell properties, lysis efficiency, genome characteristics and reference availability **[Reska et al. 2024, Bell et al. 2021, Schmidt et al. 2022]**. Large seasonal inputs of plant DNA, likely driven in part by pollen release, could therefore reduce the relative representation of fungi, bacteria and other lower-biomass components and make them more difficult to detect. This effect may be particularly important during spring, when the weekly samples showed a pronounced increase in plant reads (**Figure 3A**). Parallel size fractionation could be used to separate pollen-rich and smaller-particle fractions without discarding information from either.

Despite this potential masking effect, fungal pathogens and saprotrophs, including taxa of allergenic importance, were detected. Monitoring airborne fungal pathogens, as previously proposed **[Peers et al. 2024]**, could support surveillance of diseases such as ash dieback and assessment of the associated ecosystem impacts resulting from the loss of European ash **[Hultberg et al. 2020]**. Similarly, chytridiomycosis has spread from Asia and threatens amphibian species worldwide **[O’Hanlon et al. 2018]**, while its detection in rainwater suggests that airborne dispersal is possible **[Kolby et al. 2015]**. More broadly, the COVID-19 pandemic highlighted the importance of understanding pathogen transmission routes and monitoring genetic change in emerging pathogens. Environmental genomic surveillance could therefore provide useful information to support public-health responses during epidemics **[Yousif et al. 2023]**.

Previous eDNA studies have noted that analysis is constrained by imperfect sequence databases i.e. species gaps leading to either misassignment to the closest available species, assignment to a higher (and less informative) taxonomic group, or failure to classify the read at all **[Marques et al. 2021]**. Much of the current research looks at missing microbial genomes, and how metagenome assembled genomes (MAGs) can contribute **[Smith et al. 2022, Anthony et al. 2024]**. The genome of every species or strain in a database is not essential since it is possible to classify to a higher taxonomic level e.g. genus, based on sequence similarity to genomes that are available by a Lowest Common Ancestor algorithm **[Huson et al. 2007]**. However, we believe that the effect of database gaps seems more important in genomes where only a small fraction (≤5%) is coding or regulatory (constrained by selection); for many higher eukaryotes the vast majority of their large genome is subject to the neutral theory of molecular evolution **[Kimura 1968]** and over time becomes species specific. Indeed the vast majority of differences between human and chimpanzee genomes (4-5 million years separation) are in non-coding sequences **[Franchini and Pollard 2017]**. With greater evolutionary divergence (∼75 million years), much of the human and mouse genomes can no longer be aligned, although orthologous protein-coding sequences, which comprise only ∼1–2% of the genome, retain high sequence identity **[MGSC 2002]**.

The NHM Wildlife Garden data illustrate how reference availability interacts with biological source strength. Of the 349 catalogued garden plant species, only 74 had genome assemblies available in NCBI; 30 of these were detected, compared with only eight detections among the 275 species without genome assemblies (**Figure 5C**). However, 44 genome-represented species were still not recovered, indicating that reference availability alone did not explain detection. Detection was biased towards larger plants, with 18 of 23 large woody plants or climbers detected, compared with 5 of 20 medium herbaceous species and 2 of 21 small or low-growing herbs. Thus, airborne detection appears to reflect both database completeness and source strength, likely influenced by plant size, dispersal biology, seasonal material release and distance from the collector (**Figure 5D**).

Thus shotgun metagenomics is differently affected by lack of reference genomes and difficulty analysing species specific sequences than is metabarcoding, as such targeted sequencing can generate novel OTUs or ASVs by read clustering **[Serite et al. 2023]**. The closest to this for shotgun data is the generation of MAGs, which are less prevalent and technically harder to generate for eukaryotes **[Boulton et al. 2025]**. In our analysis we see large numbers of unassigned shotgun reads, probably due to the uniqueness of these sequences and database gaps among higher eukaryotes, but we also see reads that are assigned to *Thuja sutchuenensis* (an endangered species in the mountains of Chengkou, China) that is not reported by NBN Atlas to be present in the UK **[NBN 2026]**. These reads are likely from common *Thuja* species without reference genomes. Fortunately projects under the auspices of the Earth Biogenome Project are creating publicly available reference genomes for all eukaryotes filling such gaps with high quality genome sequences **[Blaxter et al. 2025]**, which is prioritising genome assembly e.g. choosing one representative per family or genera. Targeting plants as the commonest primary producers with outsized effects on defining food webs (pathogens, saprophytes, herbivores and predators etc.) and ecosystems, could also be a good genome prioritisation strategy especially for biodiversity monitoring.

Shotgun sequencing has much richer data than metabarcoding, so can address wider questions about the full genetics of detected organisms. For example, we profiled AMR genes within urban air samples, and previously we compared fungal spore (wheat yellow rust) derived reads to the population structure of the species using single nucleotide polymorphisms **[Giolai et al. 2024]**. Recently [**Nousias et al. 2025**] called wider polymorphism types and haplogroups from human DNA in air, and used mitogenomes to place organisms within a tree of species isolates. There are reported differences in mitochondrial and nuclear DNA decay rates which suggest that a mtDNA focus e.g. metazoan COI barcode, underestimates ecosystem composition compared to a shotgun approach **[McCauley et al. 2023]**, if this is also true for the plant plastid genome this would affect the barcode genes matK and rbcL.

## 5. Conclusion

Airborne DNA remains an underexplored source of environmental information **[Irwin 2026]**. Across London and a year of weekly sampling, our shotgun-metagenomic workflow detected diverse taxa and revealed pronounced spatial and seasonal changes in airborne communities. By combining real-time nanopore sequencing with progressive bioinformatic analysis, the workflow can generate results in under three hours while providing sequence information from across genomes rather than individual marker loci. As reference databases expand, datasets such as these can be reanalysed to classify additional reads and improve taxonomic resolution.

## Acknowledgements

This material is based upon sampling work supported by the Defense Advanced Research Projects Agency (DARPA) under Contract No. HR001119C0031. The author(s) acknowledge the support of the Biotechnology and Biological Sciences Research Council (BBSRC), part of UK Research and Innovation; Earlham Institute Strategic Programme Grant Decoding Biodiversity BBX011089/1 and its constituent work packages - BBS/E/ER/230002A (Decode WP1 Development of Novel Experimental and Bioinformatic Tools for Genomic Diversity and Analysis), BBS/E/ER/230002C (Decode WP3 Linking Fine-Scale Microbial Diversity to Ecosystem Functions). We thank Liliya Serazetdinova, Robyn Fryer and Harry Rousham for their project support during the DARPA program, and Filipa Sampaio at NHM for assistance with sample collection and processing. We thank Tom McCarter, Sylvia Myers and Fred Rumsey for providing the wildlife garden plant catalogue data.

## 6. Competing Interests / Patent Disclosure

MDC, RML, SM, MA, RM are named inventors on EU Patent No. EP4460830B1. MDC, RML, DH, DC, PA, PC, RM, MG are named inventors on EU Patent Application No. EP4562140A1, which is currently pending. Both relate to aspects of the technology described in this article and are owned and filed by Earlham Enterprises Ltd and Natural History Museum Trading Co Ltd. RML and MDC are co-founders and directors of Agnos Biosciences Ltd, a company which offers air metagenomics services. MDC also owns shares in Oxford Nanopore Technologies.

## Supplementary Figures

**Supplementary Figure 1:**
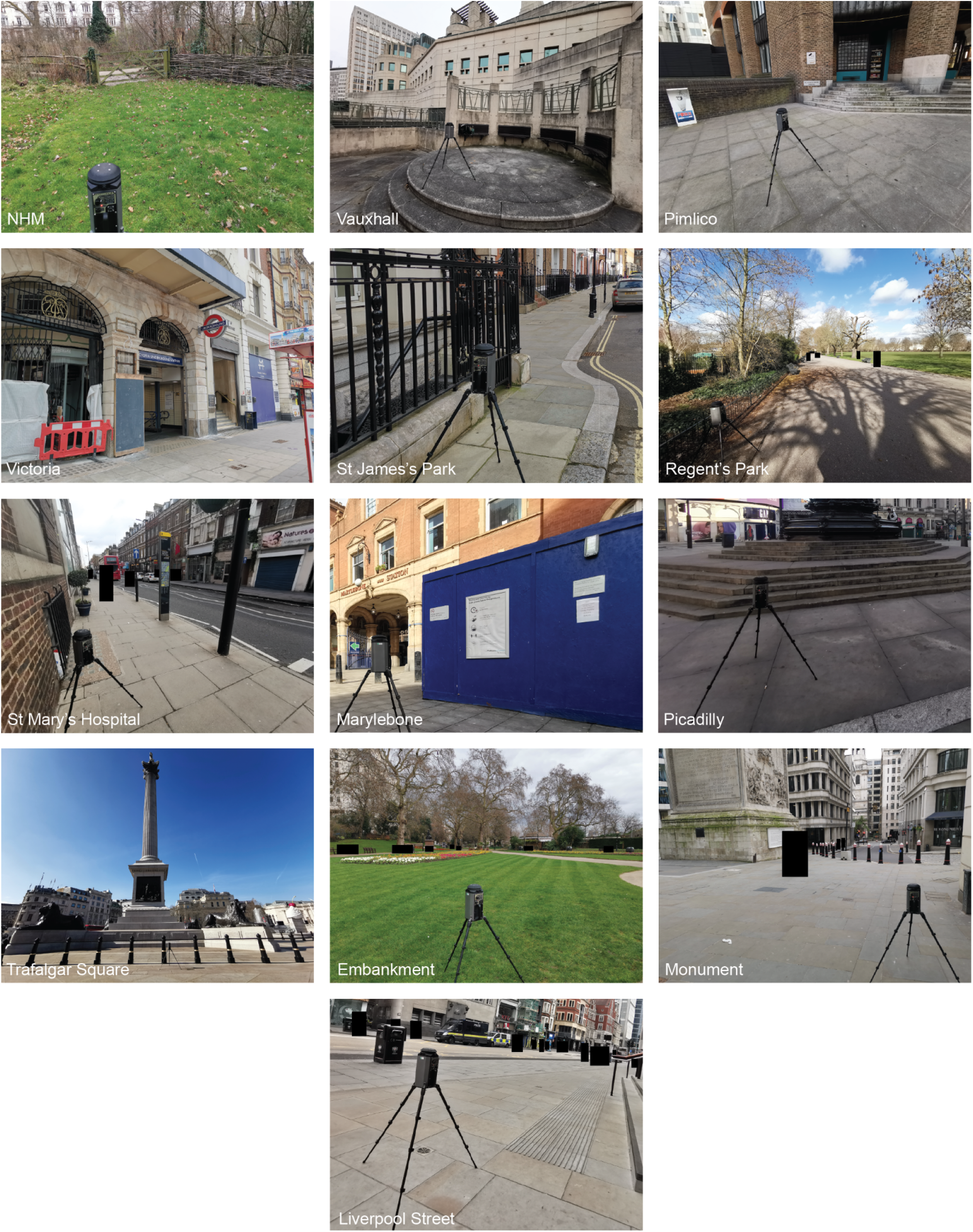
Sampling locations. Images shown are from the March 2021 sampling (LB), but the same locations were used in 2019. Pedestrians are cropped out or obscured with black rectangles.

**Supplementary Figure 2:**
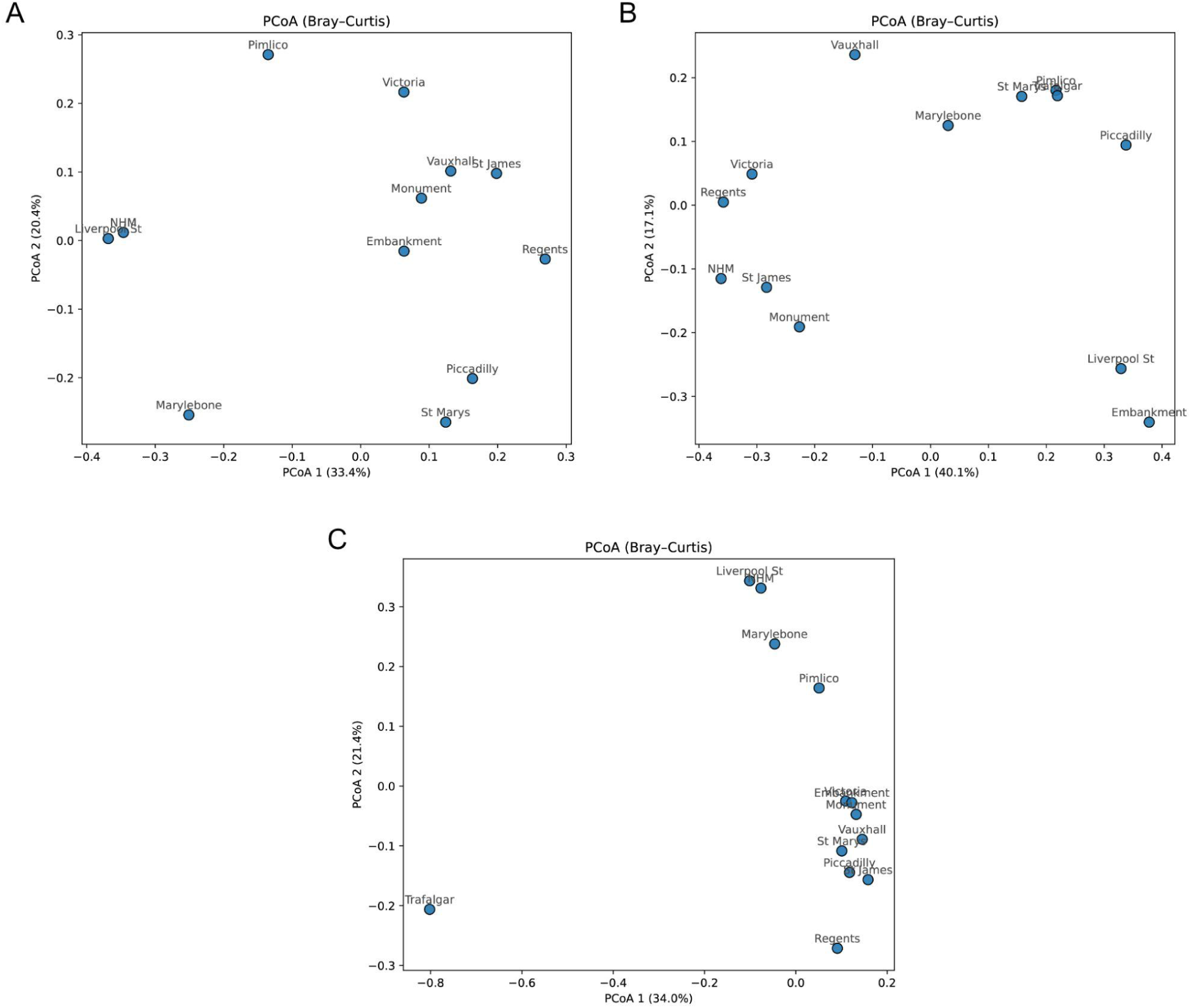
Principal Coordinate Analysis (PCoA) of the air samples at species level (≥0.005% and ≥2 reads). (A) August 2019 (LC) samples with the Trafalgar Square sample excluded due to the capture of an insect in the collector. (B) March 2021 (LB) samples. (C) August 2019 (LC) samples including Trafalgar Square.

**Supplementary Figure 3:**
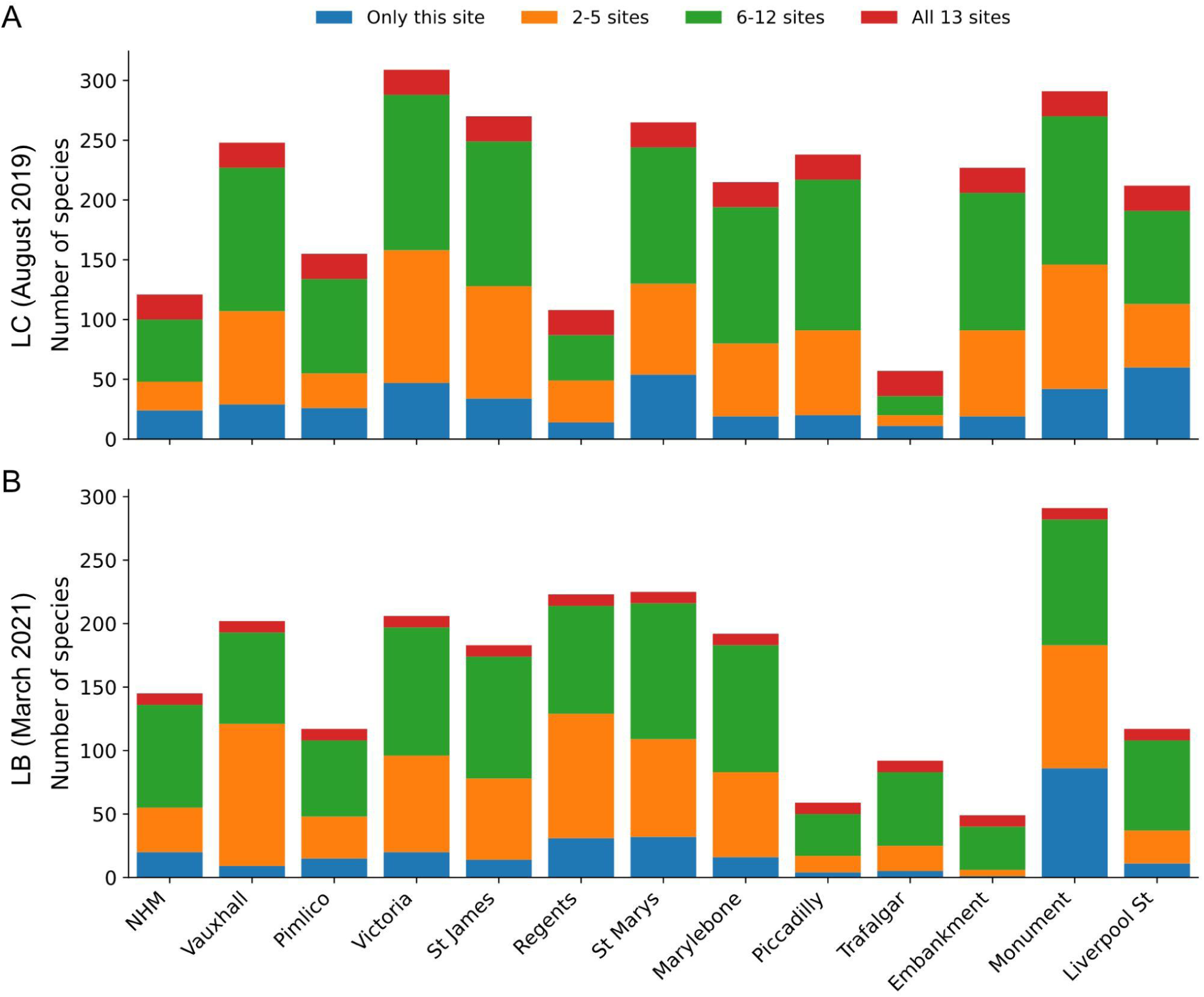
Common and unique species counts across sites (only taxa with at least 0.005% abundance considered). (A) August 2019 (LC) samples. (B) March 2021 (LB) samples.

**Supplementary Figure 4:**
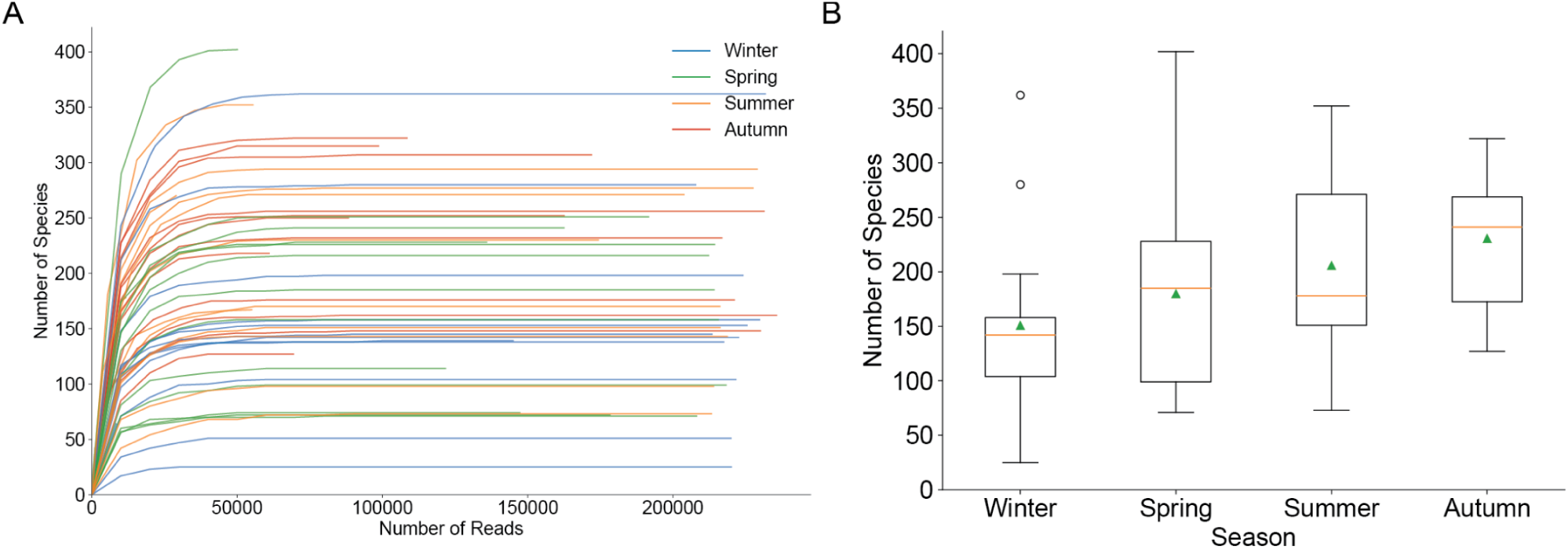
(A) Species accumulation curves for individual weekly samples, coloured by meteorological season. Curves show the cumulative number of detected species with increasing numbers of analysed reads. Species were retained if supported by ≥2 reads and representing ≥0.005% of total analysed reads in the final sample dataset. (B) Distribution of species richness per weekly sample by season. Boxes show the interquartile range with the median indicated by the central line; triangles indicate the mean, whiskers extend to 1.5 × the interquartile range, and points beyond the whiskers are outliers.

**Supplementary Figure 5:**
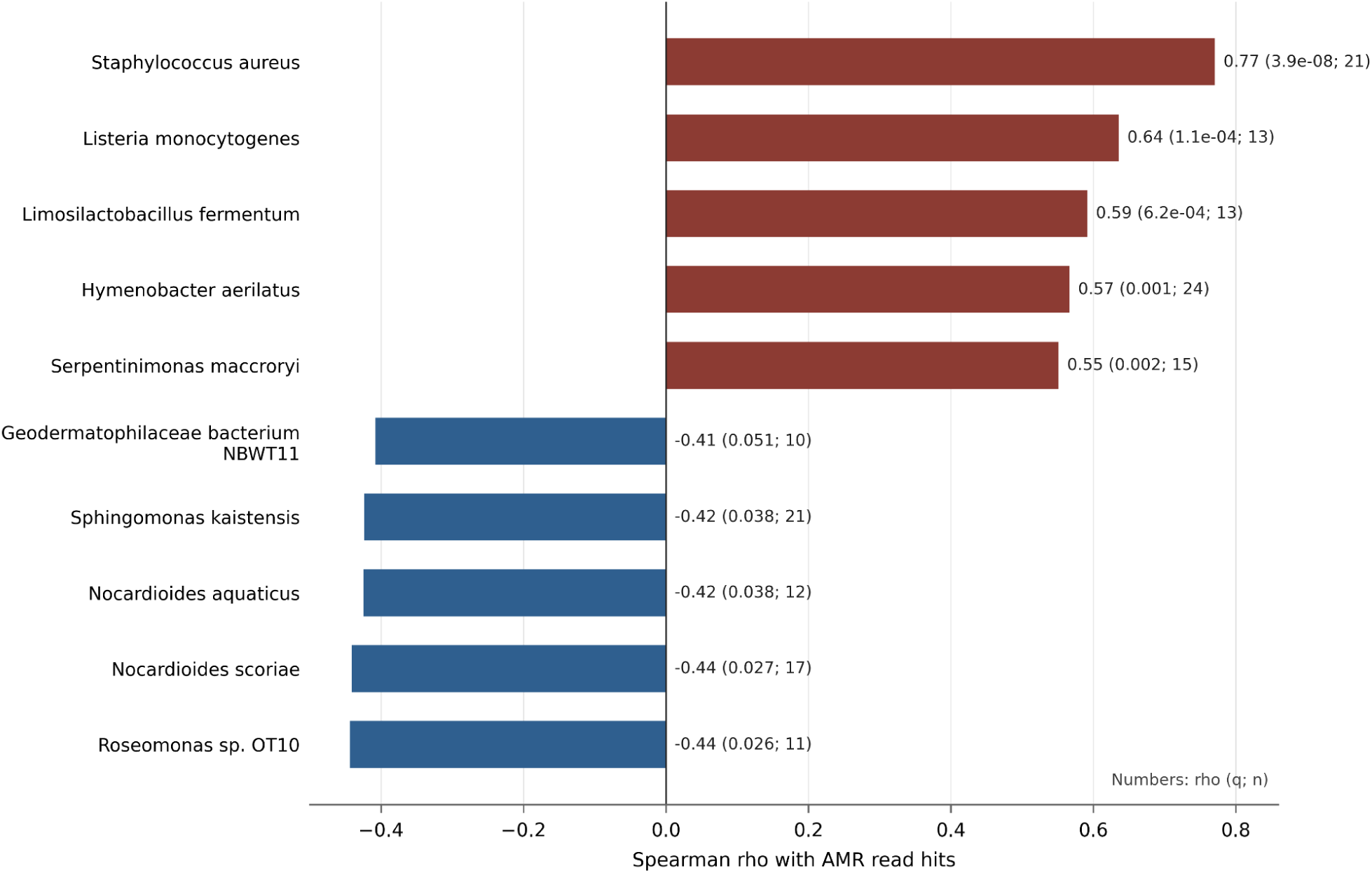
Spearman rank correlations between bacterial species relative abundance and total AMR read hits across the weekly NHM wildlife garden samples. The five strongest positive (red) and five strongest negative (blue) bacterial species associations are shown.

**Supplementary Figure 6:**
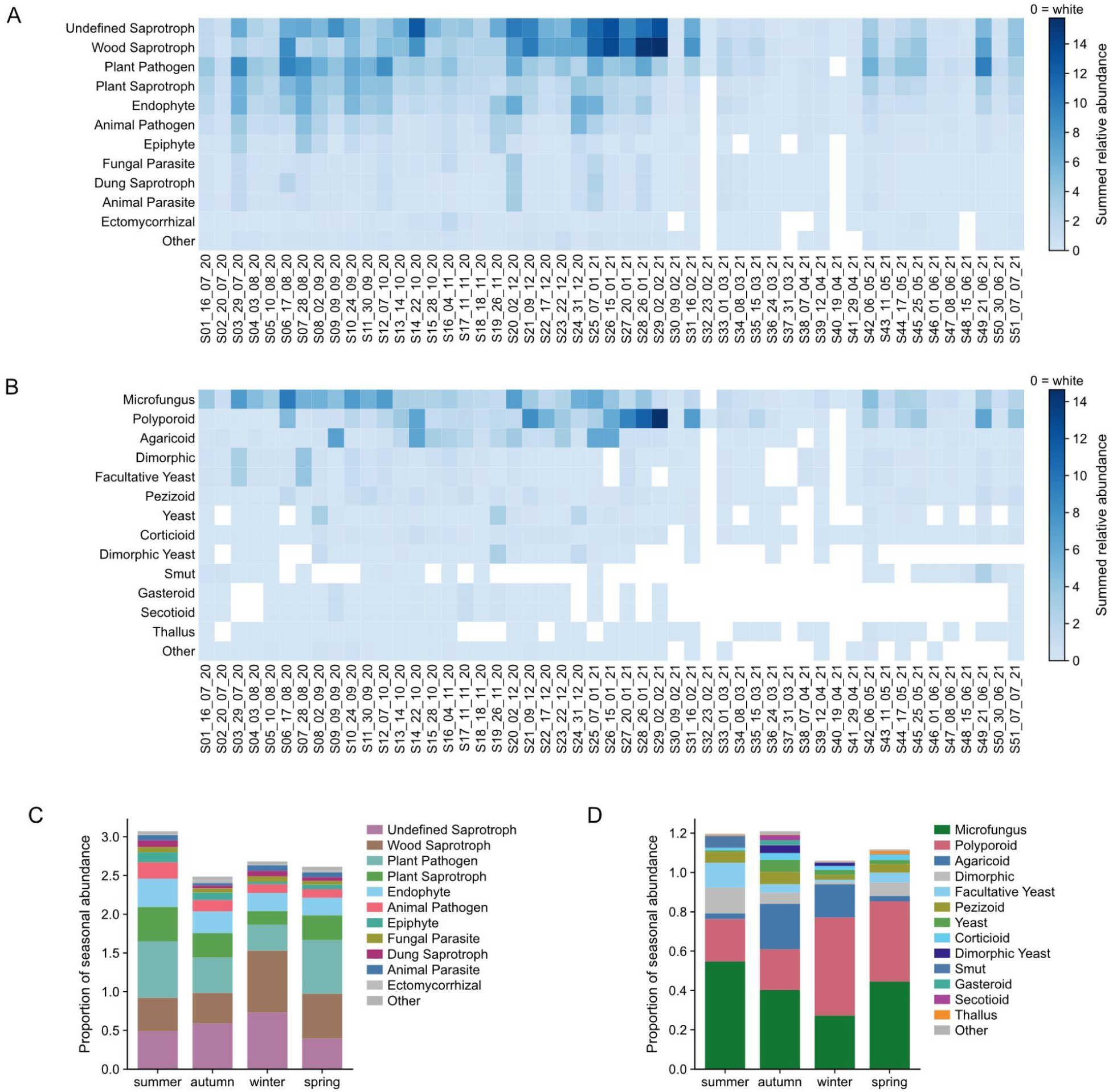
Weekly and seasonal functional composition of airborne fungi in the Natural History Museum Wildlife Garden. (A, B) Heat maps showing the summed relative abundance of fungal taxa assigned by FUNGuild to ecological guilds (A) and growth morphologies (B) across weekly samples collected from July 2020 to July 2021. Colour scales are linear and independently scaled for each heat map; white denotes zero abundance. (C, D) Stacked bars showing the normalised seasonal contributions of ecological guilds (C) and growth morphologies (D). Categories were retained when they contributed at least 1% of pooled weekly abundance or reached at least 5% in any week; remaining category contributions were combined as Other. Taxa with multiple FUNGuild assignments contributed their full abundance to each assigned category, meaning that seasonal stacked totals can exceed one.

